# A unified computational framework for modeling genome-wide nucleosome landscape

**DOI:** 10.1101/202580

**Authors:** Hu Jin, Alex I. Finnegan, Jun S. Song

**Affiliations:** Department of Physics, University of Illinois, Urbana-Champaign, Urbana, IL 61801; Carl R. Woese Institute for Genomic Biology, University of Illinois, Urbana-Champaign, Urbana, IL 61801

**Keywords:** nucleosome positioning, nucleosome occupancy, DNA sequence, statistical mechanics, cross-entropy

## Abstract

Nucleosomes form the fundamental building blocks of eukaryotic chromatin, and previous attempts to understand the principles governing their genome-wide distribution have spurred much interest and debate in biology. In particular, the precise role of DNA sequence in shaping local chromatin structure has been controversial. This paper rigorously quantifies of the contribution of hitherto-debated sequence features – including G+C content, 10.5-bp periodicity, and poly(dA:dT) tracts – to three distinct aspects of genome-wide nucleosome landscape: occupancy, translational positioning and rotational positioning. Our computational framework simultaneously learns nucleosome number and nucleosome-positioning energy from genome-wide nucleosome maps. In contrast to other previous studies, our model can predict both *in-vitro* and *in-vivo* nucleosome maps in *S. cerevisiae*. We find that although G+C content is the primary determinant of MNase-derived nucleosome occupancy, MNase digestion biases may substantially influence this GC dependence. By contrast, poly(dA:dT) tracts are seen to deter nucleosome formation, regardless of the experimental method used. We further show that the 10.5-bp nucleotide periodicity facilitates rotational but not translational positioning. Applying our method to *in-vivo* nucleosome maps demonstrates that, for a subset of genes, the regularly-spaced nucleosome arrays observed around transcription start sites can be partially recapitulated by DNA sequence alone. Finally, *in-vivo* nucleosome occupancy derived from MNase-seq experiments around transcription termination sites can be mostly explained by the genomic sequence. Implications of these results and potential extensions of the proposed computational framework are discussed

## 2. Introduction

Eukaryotic DNA is tightly packaged inside the nucleus through a hierarchical structure of chromatin. The first level of compaction involves wrapping 147 base pairs (bps) of DNA around a histone octamer, forming the nucleosome that repeatedly occurs across the genome. The precise locations of nucleosomes on genomic DNA influence higher-order chromatin structure [1] and regulates diverse cellular processes by controlling local DNA accessibility [2]. Specifically, it is believed that most DNA-binding proteins preferentially bind the unprotected linker DNA between nucleosomes. Discovering the rules that govern genome-wide nucleosome landscape is thus a major step towards understanding nucleosome-mediated chromatin structure and regulatory processes taking place on the chromatin template.

Decades of research have demonstrated that no single mechanism can explain every aspect of the nucleosome landscape, and several competing ideas have been proposed to date. First, both *in-vitro* [3, 4] and *in-vivo* [5–8] studies have suggested that DNA sequence may play a role, since experimental evidence suggests that certain sequences either favor or disfavor nucleosome formation. Second, nucleosome locations can be actively modulated by *trans*-factors, including chromatin remodelers, transcription factors, and RNA polymerase [2]. Finally, statistical positioning arising from a sharp barrier and the steric exclusion between neighboring nucleosomes may account for a large portion of nucleosome arrays around transcription start/termination sites (TSS/TTS) [9,10]. The relative contributions of these factors in shaping local chromatin structure remain unclear and, in fact, highly controversial [2–4,11,12]. In particular, to what extent the global nucleosome landscape is *a priori* programmed into the genomic sequence itself remains to be determined.

To date, various sequence features, such as the G+C content (*GC*), 10.5-bp nucleotide periodicity, and poly(dA:dT) tracts (*polyA*), have been shown to influence nucleosome formation; however, the genome-wide contributions of these individual features have not been completely characterized. Another impediment to understanding the determinants of genome-wide nucleosome landscape stems from the confusion about different technical concepts involved [13, 14]. That is, different sequence features may play different roles in affecting nucleosome occupancy – referring to the probability of a given genomic location being occupied by a nucleosome – versus nucleosome positioning – referring to the precise genomic locations of individual nucleosomes, which can be described by the start or center (dyad) locations of the nucleosomes [13, 14]. The distinction between these two concepts is often blurred in the literature. More precisely, the concept of nucleosome positioning can be further distinguished into translational and rotational positions. The translational nucleosome position refers to the location of the 147-bp DNA contacting the histone octamer, while the rotational nucleosome position refers to the rotational orientation of DNA double helix relative to the histone surface [13, 14]. Previous studies have focused on either a single sequence feature (e.g., [15] studies the role of 10.5-bp periodicity in genome-wide nucleosome occupancy and nucleosome positioning in synthetic DNA sequences) or a single aspect of nucleosome landscape (e.g., [16] studies the role of *GC* and 10.5-bp periodicity in nucleosome occupancy). By contrast, this paper presents a unified framework for quantitatively characterizing the role of several distinct sequence features (*GC*, 10.5-bp periodicity, and *polyA*) in nucleosome occupancy, translational nucleosome positioning, and rotational nucleosome positioning, both *in vitro* and *in vivo*. To avoid confusion in the rest of the paper, the term “nucleosome landscape” will be used to refer to the general biological problem encompassing nucleosome occupancy, translational nucleosome positioning, and rotational nucleosome positioning.

The “beads-on-a-string” structure of nucleosome-occupied DNA is reminiscent of the “Tonks gas” model [17] of non-overlapping one-dimensional (1D) rods. Indeed, this model has been used to quantify the role of statistical positioning and explain the regularly-spaced nucleosome arrays observed around TSS and TTS [9, 10, 18, 19]. In order to further account for sequence effects, a 1D sequence-dependent binding energy can be introduced into the model. This resulting model of 1D hard rods subjected to an external potential has been widely applied to study the genome-wide distribution of nucleosomes [15, 16, 20–26]. In particular, Locke *et al.* consider the inverse problem of inferring the nucleosome-positioning energy from genome-wide nucleosome maps generated through high-throughput sequencing, regressing this energy as a linear function of DNA sequence content [16]. They conclude that *GC* is the primary determinant of nucleosome occupancy, with the 10.5-bp periodicity making little contribution, and that DNA sequence alone cannot create the nucleosome arrays observed around TSS and TTS *in vivo*. The work of Locke *et al.* has provided refreshing perspectives in the field, and their approach has been extended in several subsequent studies [23, 24, 26]. However, we here show that the calculation performed by Locke *et al.* incorrectly estimates the total number of nucleosomes in the genome (we refer to this number as the nucleosome number, Materials and Methods, Section 1). We demonstrate that this problem ultimately leads to incorrect inference of the nucleosome-positioning energy, thereby confounding the downstream analysis of sequence effects (Materials and Methods, Section 1).

The main difficulty in the approach of Locke *et al.* arises from their normalization scheme for converting the raw counts of sequencing reads to well-defined probability values. In this paper, we first develop a cross-entropy method (CEM) that gets around this normalization problem and simultaneously learns nucleosome number as well as nucleosome-positioning energy from genome-wide nucleosome maps (Materials and Methods, Section 2). Our CEM is inspired by similar optimization-based approaches [15, 25, 26], but contains no artificial components (as in [15, 26]) and provides a flexible framework that can incorporate various regulatory factors, such as chromatin remodelers, which may affect nucleosome occupancy and positioning (Discussion section).

We validate our method using several independent nucleosome maps in *S. cerevisiae* and demonstrate that it consistently outperforms the method of Locke *et al.* in predicting nucleosome occupancy, benefiting from the ability to learn the correct nucleosome number, even though CEM contains much fewer free parameters (Results, Section 1). We then present a comprehensive quantification of the influence of *GC*, 10.5-bp periodicity, and *polyA* on nucleosome occupancy and positioning. Importantly, we show that the *GC* dependence of MNase-derived nucleosome occupancy is substantially skewed by MNase digestion biases [27, 28], while *polyA* excludes nucleosome occupancy regardless of the experimental method used (Results, Sections 2 & 3). We also establish that the 10.5-bp periodicity, although not significantly affecting nucleosome occupancy, facilitates rotational but not translational positioning [29] (Results, Section 4). Further application of CEM to *in-vivo* nucleosome maps shows that the oft-observed regularly spaced nucleosome array at TSS can be partially recapitulated by DNA sequence alone for a subset of genes, in sharp contrast to the conclusion of Locke *et al.* (Results section 5, 6). We argue that this discrepancy stems from the previous under-estimation of nucleosome number. Finally, we show that MNase-derived nucleosome occupancy around TTS in *S.cerevisiae* is mostly encoded in the genomic sequence (Results, Section 7). Our work thus rigorously quantifies of the effect of DNA sequence on nucleosome formation and highlights the importance of correctly estimating nucleosome number in modeling *in-vivo* and *in-vitro* nucleosome landscapes.

## 3. Materials and Methods

### 3.1 Evaluation of the effect of estimated nucleosome number on model predictions

Nucleosome distribution can be modeled as the hard-core Tonks gas of 1D rods [16, 23, 24, 30] (Supplementary Methods, Section 1.1). In the simplest form, nucleosomes are assumed to occupy a fixed number *a* of base pairs (bps) along DNA of length *L* and cannot overlap with each other due to steric exclusion. This steric exclusion is realized by imposing hard-core interaction between neighboring nucleosomes. The total energy of the system then involves only the sequence-dependent nucleosome-positioning energy *u* due to histone-DNA interactions, with each nucleosome occupying bps *i*, *i* + 1,…, *i*+*a*−1 contributing an energy term *u*(*i*) at location *i*. Given a fixed form of *u*, the statistical mechanics of nucleosome landscape concerns solving for the one-particle distribution function *n*_1_(*i*), the probability of a nucleosome starting at bp *i*, and the nucleosome occupancy 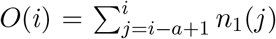, the probability of bp *i* being occupied by some nucleosome. This problem of calculating *n*_1_ and *O* from known *u* is termed the direct problem, which can be solved using techniques from statistical mechanics. However, the inverse problem of calculating *u* from known *n*_1_ and *O* is more difficult and also more pertinent in biology, since *n*_1_ and *O* can be directly linked to experimental data, as described below.

This 1D hard-rod model and its variants have been widely applied to study various questions associated with genome-wide nucleosome landscapes [9, 10, 15, 16, 20–26]. In particular, Locke *et al.* have used this approach to investigate how DNA sequence regulates nucleosome occupancy [16]. Specifically, *n*_1_ and *O* are first obtained from high-throughput sequencing data of genome-wide nucleosomes; then, the nucleosome-positioning energy *u* (or rather, 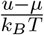, where *μ* denotes chemical potential, *k*_*B*_ the Boltzmann constant, and *T* temperature) is calculated by solving the inverse problem in the Grand Canonical Ensemble. To simplify notation, we use *u* to denote the dimensionless quantity 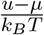 throughout the paper. Finally, linear models on DNA sequences are fitted to the calculated *u* to study how different sequence features influence the nucleosome energetics (Supplementary Methods, Section 1.2). We here describe an important problem associated with their very first step of estimating *n*_1_ and *O* from experimental data and demonstrate how this problem might lead to incorrect conclusions about the sequence dependence of nucleosome landscape.

Raw data from a high-throughput genome-wide nucleosome mapping experiment, such as MNase-seq, yield the nucleosome count *ñ*_1_(*i*) and the observed nucleosome occupancy 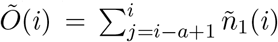, where *ñ*_1_(*i*) is the number of sequenced nucleosomes starting at bp *i* and *Õ*(*i*) is the the number of sequenced nucleosomes covering bp *i*. Under the assumption that the true one-particle distribution function is proportional to the observed nucleosome count, i.e. *n*_1_ ∝ ñ_1_, *n*_1_ may be estimated by properly scaling *ñ*_1_. In Locke *et al.*, *n*_1_ is estimated as

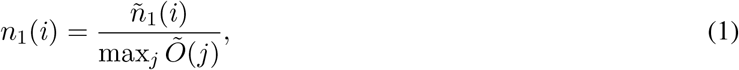

such that the obtained *n*_1_ and *O* are both bounded between 0 and 1 and, thus, have probabilistic interpretations.

We note several problems associated with this estimation. First, it artificially sets the maximum value of *O* to 1; in other words, at least one bp in the genome is assumed to have probability 1 of being occupied by some nucleosome, but this assumption is not necessarily true. Second, the normalization coefficient max_*j*_ *Õ*(*j*) is determined by read counts in only a very small region of the genome and, thus, may be sensitive to experimental noise and bias. Most importantly, Equation 1 does not guarantee the correct estimation of nucleosome number. In the Grand Canonical Ensemble, the ensemble-averaged nucleosome number in the entire genome is (Supplementary Methods, Section 1.1.3),

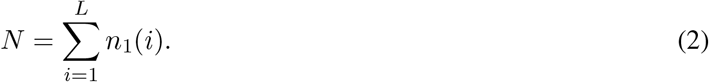

If *N* were known, the theoretically correct estimate of *n*_1_ would be

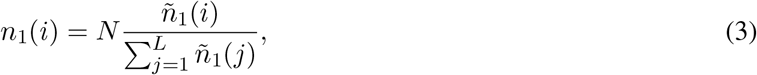

such that Equation 2 is automatically satisfied. Since 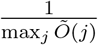 is not necessarily equal to 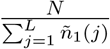, however, the normalization in Equation 1 does not guarantee the condition in Equation 2. In fact, the nucleosome number estimated by Equation 1 is solely determined by the total sequencing depth and the denominator max_*j*_ *Õ*(*j*), which may be sensitive to experimental noise and bias.

The incorrect estimation of nucleosome number using Equation 1 may confound the inference of nucleosome-positioning energy *u* and the subsequent analysis of its sequence dependence. We illustrate this issue using a simple simulation, where the nucleosome-positioning energy is fixed to be proportional to the G+C content (*GC*) of occupied nucleosomal sequence; that is, 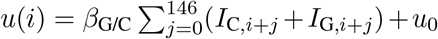, where β_G/C_ = −0.2, *u*_0_ = 10, and *I*_*α,i*_ is the indicator function for nucleotide *α* at location *i*. Figure 1 shows the simulation results using the genomic sequence on chromosome I (chrI) of *S. cerevisiae*. Since energy *u*(*i*) is completely determined by the DNA sequence and known, *n*_1_ and *O* can be calculated by solving the direct problem (Figure 1A). The nucleosome number on chrI is then calculated to be *N* = *∑*_*i*_*n*_1_(*i*)= 1276. An array of well-positioned nucleosomes is evident between the two energy barriers at the locus shown in Figure 1A, reminiscent of statistical positioning [9, 10, 18, 19]. Conversely, assuming that the true *n*_1_ and *O* are known, energy *u* can be calculated by solving the inverse problem (Figure 1B). We then fit a linear model of energy that depends only on *GC* of nucleosome-occupied sequence. As expected, the learned parameters are exactly the same as the true values, 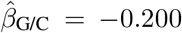 and *û*_0_ = 10.0, with *R*^2^ = 1.00; consequently, the predicted 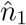 and *Ô* obtained from this estimated energy function reproduces the true *n*_1_ and *O* (Figure 1B). Next, to simulate the incorrect estimation of nucleosome number by Equation 1, we have scaled *n*_1_ down globally such that the estimated nucleosome number is 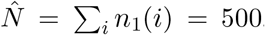. The energy calculated from this incorrectly normalized *n*_1_ significantly deviates from the true energy (Figure 1C; c.f. Figure 1A). In this case, a linear model depending only on *GC* cannot fully capture the incorrectly learned energy (*R*^2^ = 0.186), and the fitted parameters, 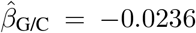 and *û*_0_ = 6.79, are also off from the true values. Furthermore, the 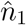 and *Ô* profiles predicted from the fitted linear energy function do not agree with the true *n*_1_ and *O*. In particular, the predicted profiles in Figure 1C lack a nucleosome array between the two energy barriers present in the true data shown in Figure 1A. Therefore, incorrect normalization of *n*_1_ can easily lead to incorrect inference of nucleosome-positioning energy and incorrect predictions of nucleosome profiles; and, conclusions about the sequence-dependence of nucleosome landscape drawn from analyses using this normalization scheme may be confounded.

**Figure 1:**
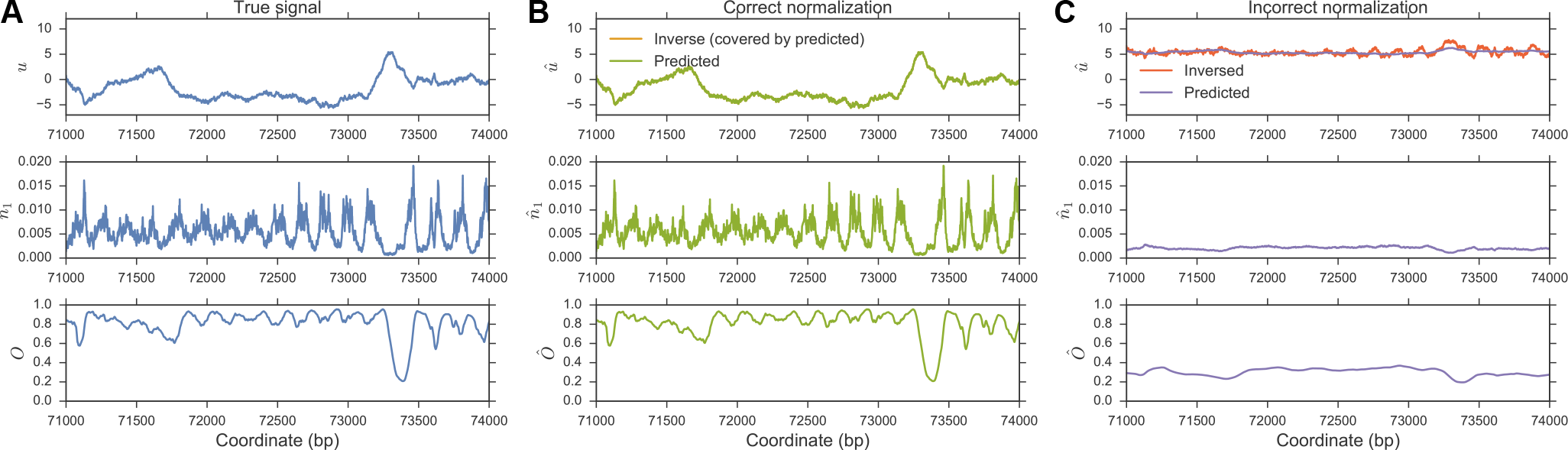
Incorrect estimation of nucleosome number may distort the inference of nucleosome-positioning energy. (A) True energy (top panel), *n*_1_ (middle panel), and occupancy *O* (bottom panel) in the simulation. (B) Using the true *n*_1_ and *O* from (A), nucleosome-positioning energy is calculated by solving the inverse problem (orange curve in top panel, hidden by the overlapping green curve). By fitting this calculated energy to a linear sequence model that depends only on *GC*, the positioning energy can be predicted based on sequence (green curve in top panel). 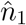 (middle panel) and *Ô* (bottom panel) are predictions from this fitted linear energy model. (C) Same as (B) except that the true *n*_1_ is scaled down, so that the nucleosome number is now only 500 (see main text).

It is important to note that the above simulation is not an intentional exaggeration of the potential problem. On the contrary, according our investigation, using Equation 1 on various *in-vivo* nucleosome maps in *S. cerevisiae* yields an estimated nucleosome number of at most ~ 30000 (Supplementary Table S1), whereas, as shown in the Results section, the correctly estimated nucleosome number is more than 60000, which is close to the common belief that about 80% of the yeast genome is occupied by nucleosomes *in vivo* [31]. Thus, the normalization in Equation 1 underestimates the nucleosome number by half in real *in-vivo* data sets, close to the scenario depicted in Figure 1.

Finally, we point out that Equation 3, despite being justified as a maximum likelihood estimate, has its own problems. First, the precise nucleosome number *N* is unknown, and estimating *N* would require some nucleosome calling algorithm which usually depends on an arbitrary choice of thresholds. Another issue is that the estimated *n*_1_ and *O* are not guaranteed to be between 0 and 1, leading to difficulties in downstream analysis, such as solving the inverse problem to obtain *u*. To avoid these issues, we will develop in the following section a method that learns *N* and *u* simultaneously from experimental data, without requiring a direct estimate of *n*_1_ beforehand.

### 3.2 A cross-entropy method for simultaneously learning nucleosome number and nucleosome-positioning energy

The above difficulty faced by the Locke method (LM) arises because solving the inverse problem of inferring the nucleosome-positioning energy *u* requires estimating *n*_1_ and *O* as the first step. In order to get around this difficulty, we avoid solving the inverse problem and instead learn *u* through solving the following optimization problem (Supplementary Methods, Section 1.3),

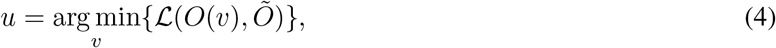

where *Õ* denotes the observed nucleosome occupancy from raw data as previously defined, *O*(*υ*) the nucleosome occupancy predicted from nucleosome-positioning energy *υ*, and 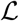 a loss function that measures the difference between prediction *O*(*υ*) and observed *Õ*. Throughout the paper, we use a weighted version of negative Pearson correlation coefficients as the loss function (see below). The obtained nucleosome-positioning energy *u* thus predicts a nucleosome occupancy profile that best correlates with the observed one. Since the Pearson correlation coefficient is invariant under linear transformations of its arguments, no scaling or normalization of *Õ* is needed beforehand, and the difficulty described in the previous section thus completely disappears. Importantly, the information of nucleosome number is implicitly contained in the overall shape of *Õ*, and our method is able to automatically adjust the nucleosome number to match the observed nucleosome occupancy profile. We will show in the Results section that this method can indeed learn the correct nucleosome number empirically observed in high-throughput nucleosome maps. Since the Pearson correlation coefficient can be inflated by extreme outliers, we filter out outlier regions based on observed nucleosome occupancy, resulting in a disjoint set of “good regions” along the genome (Supplementary Methods, Section 1.3). Average Pearson correlation coefficient across these good regions weighted by the region lengths is then used as the final loss function (Supplementary Methods, Section 1.3).

Naively solving the optimization problem in Equation 4 is prohibitive, since the number of parameters is of order of the genome length, with *u*(*i*) at each genomic location *i* being a free parameter. In order to reduce the parameter space, we enforce a linear sequence-dependent model:

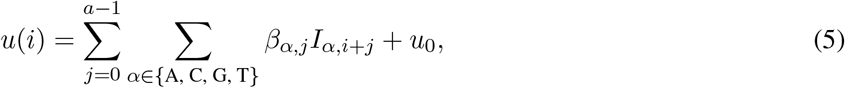

where *I*_*α*,*i*+*j*_ again denotes the indicator function of nucleotide *α* at genomic location *i*+*j*, *β*_*α,j*_ the linear coefficient characterizing how much nucleotide *α* at position *j* relative to the start of nucleosome contributes to energy *u*(*i*), and *u*_0_ the sequence-independent offset. Note that this paper focuses on mono-nucleotide models, as in Equation 5, but a generalization to *k*-mer models with *k* > 1 is straightforward. Using this linear sequence-dependent model, the number of free parameters is only several hundreds when *a* is set to the canonical nucleosome size 147.

To further reduce the number of parameters, we use a template-based model for the linear coefficient *β*_*α*,*j*_,

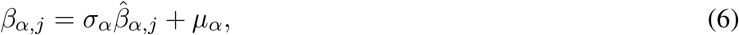

where 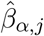 denotes a pre-defined template of nucleotide *α*, and *σ_α_* and *μ_α_* are free parameters modulating the standard deviation and mean of the template, respectively. The template 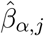 may represent our prior belief about how much different nucleotides at different relative positions contribute to the nucleosome-positioning energy. We construct this template from the negative average nucleotide frequency of nucleosomal sequences obtained from raw sequencing data (Supplementary Methods, Section 1.3). Briefly, let 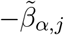 denote the average frequency of nucleotide *α* at position *j* relative to the start of the nucleosome, calculated from identified nucleosomes. The larger the value of 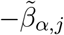, the stronger the nucleosome preference for nucleotide *α* at relative position *j*; thus, nucleotide *α* at *j* should empirically contribute more negative energy, justifying our choice. For each nucleotide *α*, we then standardize 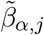 to obtain the template 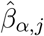 that has mean 0 and standard deviation 1 with respect to *j*. During optimization, *σ*_*α*_ in Equation 6 are constrained to be nonnegative.

Note that simple nucleosome calling algorithms can be used to obtain 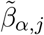 from the raw sequencing data, without worrying about estimating the nucleosome number accurately, since we only need the average nucleotide frequency (Supplementary Methods, Section 1.3). Furthermore, we enforce reverse-complement symmetry in the model by requiring 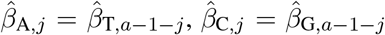, *σ*_A_ = *σ*_T_ := *σ*_A/T_, *σ*_C_ = *σ*_G_ := *σ*_C/G_, *μ*_A_ = *μ*_T_ := *μ*_A/T_, and *μ*_c_ = *μ*_G_ := *μ*_C/G_ (Supplementary Methods, Section 1.3), such that the energy *u*(*i*) calculated from any nucleotide sequence is the same as that calculated from its reverse-complement. Finally, it is easy to see that only one of *μ*_A/T_ and *μ*_C/G_ is independent, due to the constraint Σ_*α*∈{A C G T}_ *I*_*α*,*j*_ = 1, ∀*j*(Supplementary Methods, Section 1.3), allowing us to set μ_A/T_ = 0. The resulting sequence-dependent energy model is thus

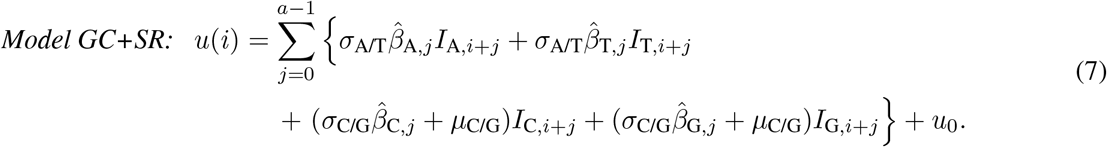

The motivation behind the choice of model name is as follows. Note that there are in total only four free parameters in the model above: *σ*_A/T_, *σ*_C/G_, *μ*_C/G_, and *u*_0_. Solving the optimization problem in Equation 4 amounts to finding the optimal values of these four parameters, such that the predicted occupancy has the highest correlation with the observed one. *σ*_A/T_ and *σ*_C/G_ characterize how the spatially resolved (*SR*, as dubbed in Locke *et al.*) pattern of the template 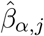 modulates the positioning energy; similarly, *μ*_C/G_ captures the energy dependence on G+C content (*GC*) of the underlying sequence, thus the name *Model GC+SR*. *u*_0_ effectively modulates the chemical potential of the system and, thus, the nucleosome number (recall that *u* denotes 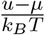 throughout the paper).

In order to study how different sequence features, especially *GC*, 10.5-bp periodicity, and *polyA*, regulate nucleosome positioning, we constructed two other models with different complexities: *Model GC*, where the energy depends on *GC* only, given by

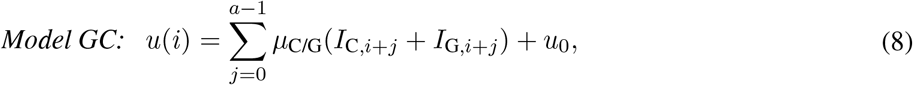

and *Model GC+SR+polyA*, where all three components contribute to the energy, given by

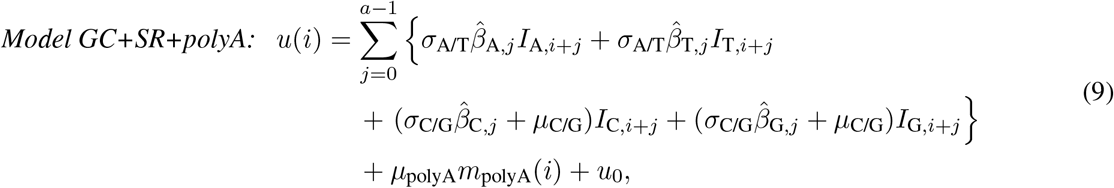

where *m*_polyA_(*i*) denotes the number of bps belonging to a poly(dA:dT) tract (defined as a homopolymer of A or T of length ≥ 5) from *i* to *i* − *a* + 1, and *μ*_polyA_ characterizes the energy dependence on poly(dA:dT) content. Note that the 10.5-bp periodicity is characterized as part of the spatially resolved sequence motif in 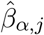. The three models *Model GC*, *Model GC+SR*, and *Model GC+SR+polyA* constitute a hierarchy of sequence models, along the same line as in Locke *et al.*, allowing us to dissect the contributions from *GC*, *SR*, and *polyA* to nucleosome positioning, respectively.

To optimize the cost function, we apply the cross-entropy method [32–35] (Supplementary Methods, Section 1.3). We note that similar ideas of learning the nucleosome energetics through optimization have been used in previous studies [15, 25, 26]. However, [15] mainly focuses on how dinucleotide periodicity affects nucleosome positions in synthetic DNA sequences and genome-wide nucleosome occupancy, and it does not analyze other sequence features such as *GC* and *polyA*; it also does not address genome-wide nucleosome positioning; finally, the proposed energy model uses artificial sinusoidal functions. Both [25] and [26] mostly focus on the effect of nucleosome unwrapping. In particular, [25] does not study the sequence-dependence of nucleosome occupancy and positioning. The sequence-dependent nucleosome-positioning energy in [26] is obtained by solving the inverse problem as in Locke *et al.* and thus has the same normalization problem described above. By contrast, our proposed method involves no artificially constructed components and provides a flexible framework that can incorporate several distinct biological features in the energy model, as illustrated by the hierarchical models in Equations 7, 8, and 9. Most importantly, our method automatically learns the nucleosome number together with the nucleosome-positioning energy, as will be demonstrated in the following Results sections.

### 3.3 Additional information on datasets and computational details

A list of datasets analyzed and their respective labels is provided in Supplementary Methods, Section 1.4. Supplementary Methods also contains an additional description of computational analyses, including the sequencing data analysis (Supplementary Methods, Section 1.5) and partial correlation analysis (Supplementary Methods, Section 1.6).

## 4 Results

### 4.1 The cross-entropy method correctly learns the nucleosome number and improves the prediction of nucleosome occupancy

We applied the cross-entropy method (CEM) (Materials and Methods; Supplementary Methods, Section 1.3) to three independent *in-vitro* nucleosome maps in *S. cerevisiae* (Supplementary Methods, Section 1.4): two using salt dialysis followed by MNase-seq (*Kaplan-MNase-invitro-salt* [3] and *Zhang-MNase-invitro-salt* [4]) and another using AcF assembly followed by MNase-seq (*Zhang-MNase-invitro-ACF* [4]). ACF is an ATP-dependent chromatin assembly factor in *Drosophila melanogaster*. *Model GC*+*SR* (Equation 7) was trained on the three datasets separately, with the templates 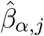 calculated from identified nucleosomes in each dataset. From the learned parameters (Supplementary Table S2), we then predicted nucleosome occupancy *O* and nucleosome number *N* = ∑_*i*_ *n*_1_(*i*) (Table 1) by solving the direct problem (Materials and Methods; Supplementary Methods, Section 1.1). To compare with Locke *et al.*’s method (LM) [16], we first estimated *n*_1_ via Equation 1, calculated the nucleosome-positioning energy *u* by solving the inverse problem (Materials and Methods; Supplementary Methods, Section 1.1), fitted a linear model as in Equation 5 (Materials and Methods; Supplementary Methods, Section 1.2), and finally predicted nucleosome occupancy *O* and nucleosome number *N* (Table 1) using the fitted sequence model.

Since both calculations using CEM and LM accounted for contributions from *GC* and *SR*, a difference in their performance is not attributable to different sequence features used by these approaches. Nucleosome occupancy predicted by both methods highly correlated with the observed one (Table 1). However, CEM achieved better correlation in all three datasets, despite the fact that the LM linear model (Equation 5) contains many more free parameters than the CEM *Model GC*+*SR* (Equation 7). In particular, CEM improved the correlation in *Zhang-MNase-invitro-ACF* the most, by about 0.1. Interestingly, the predicted nucleosome number of LM differs from that of CEM the most in the same dataset (Table 1), indicating that an incorrectly estimated nucleosome number might influence the performance of LM (Materials and Methods). Indeed, by simply tuning the nucleosome number (or equivalently, *u*_0_ in Equation 5), while keeping the learned energy from LM unchanged, we could achieve better correlations between the predicted and observed nucleosome occupancy in all three datasets (Supplementary Figure S1). By contrast, the solutions found by CEM already achieved the maximum correlations (Supplementary Figure S1). Moreover, the maximum correlations obtained by tuning the nucleosome number in LM models were still lower than the correlations achieved by CEM (Supplementary Figure S1). These results suggest that CEM is indeed learning the correct nucleosome number and demonstrate that it improves the prediction of nucleosome occupancy compared to LM.

**Table 1::**
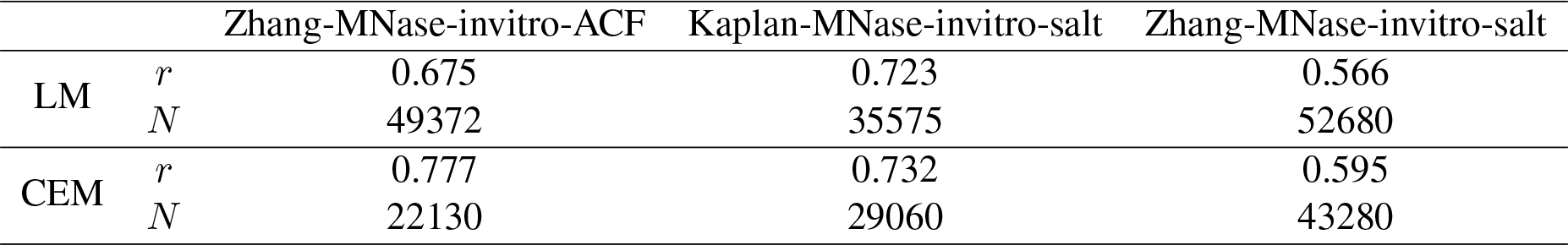
Predicted nucleosome number (*N*) and Pearson correlation coefficient (*r*) between observed and predicted nucleosome occupancy on three *in-vitro* datasets. As previously described, *r* was calculated within each “good region” after filtering out outlier regions (Supplementary Methods, Section 1.5). Table shows the average *r* weighted by region lengths.

We plotted the predicted and observed nucleosome occupancy at the GAL10-1 locus (Figure 2) and also aligned them at TSS/TTS (Supplementary Figure S2) for all three *in-vitro* datasets analyzed. Since larger nucleosome number means more tightly packed nucleosomes, we could empirically order the nucleosome numbers in these three datasets from the overall shape of the observed nucleosome occupancy (black curves in Figure 2 and Supplementary Figure S2). Specifically, *Zhang-MNase-invitro-salt* showed relatively tightly packed nucleosome arrays within the gene bodies of GAL10 and GAL1 (black curve in Figure 2C) and around TSS/TTS (black curve in Supplementary Figure S2E, F), while the nucleosome arrays were less pronounced in *Kaplan-MNase-invitro-salt* (black curves in Figure 2B and Supplementary Figure S2C, D) and hardly visible in *Zhang-MNase-invitro-ACF* (black curves in Figure 2A and Supplementary Figure S2A, B). These observations thus suggested that the nucleosome number was the largest in *Zhang-MNase-invitro-salt*, modest in *Kaplan-MNase-invitro-salt*, and the smallest in *Zhang-MNase-invitro-ACF*. Indeed, a quantitative estimation of nucleosome number by comparing the Fourier transform of predicted and observed nucleosome occupancies in the frequency domain confirmed these empirical arguments (Supplementary Figure S3).

**Figure 2::**
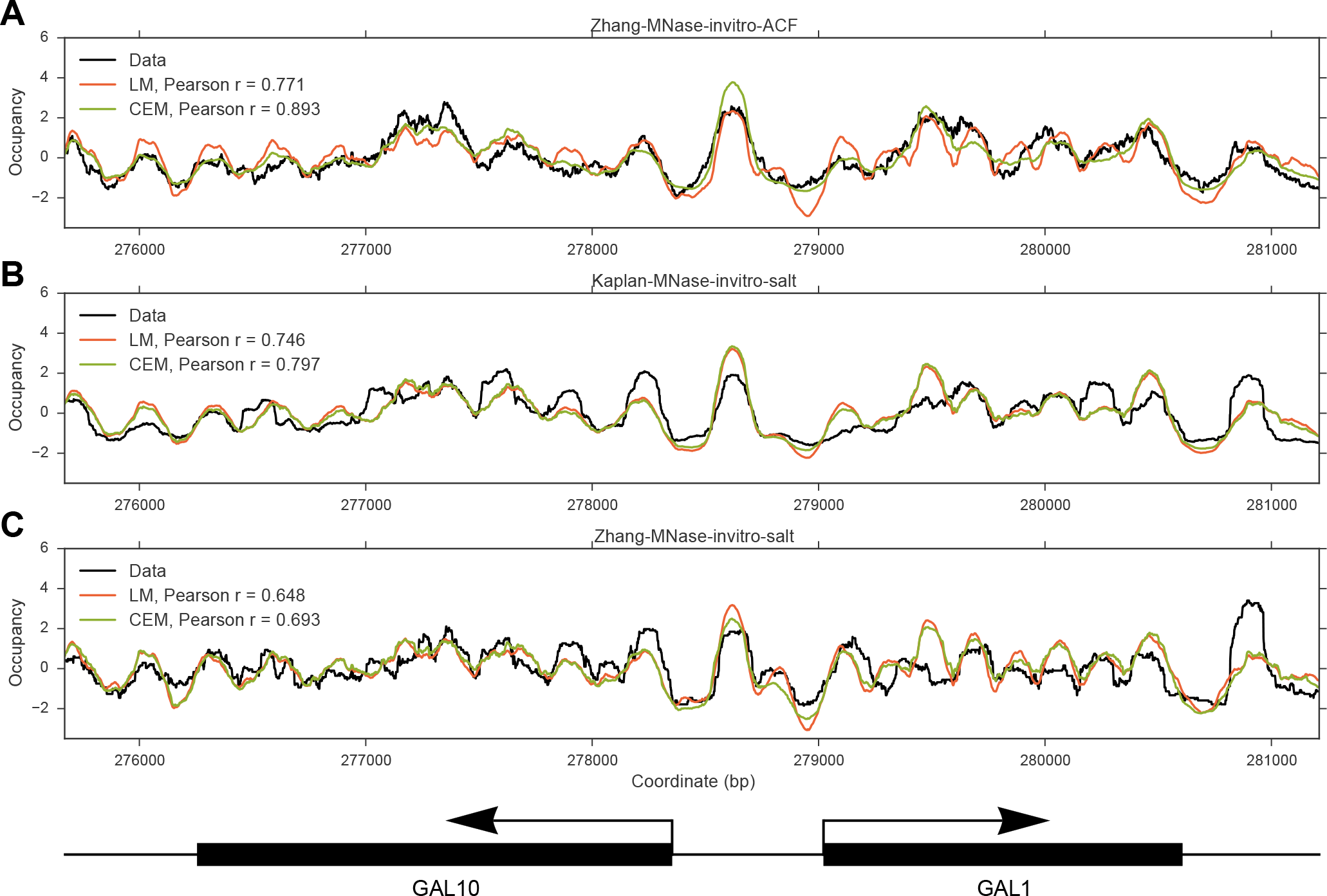
An example of a genomic locus illustrating that CEM nucleosome occupancy predictions better correlate with the observed profiles compared to LM predictions. (A) Observed (black curve), LM predicted (red curve), and CEM predicted (green curve) nucleosome occupancy for *Zhang-MNase-invitro-ACF*. (B) Same as (A), but for *Kaplan-MNase-invitro-salt*. (C) Same as (A), but for *Zhang-MNase-invitro-salt*. Each curve was standardized by subtracting the mean and then dividing by the standard deviation within the shown region.

The nucleosome numbers predicted by CEM were consistent with this ordering, while those predicted by LM were not (Table 1), further demonstrating the ability of CEM to better learn the correct nucleosome number. In particular, LM predicted a much larger nucleosome number than CEM for *Zhang-MNase-invitro-ACF*, and the predicted nucleosome number 49372 was actually closer to the number 43280 predicted by CEM for *Zhang-MNase-invitro-salt*. As a result, the nucleosome occupancy profile predicted by LM in *Zhang-MNase-invitro-ACF* (red curves in Figure 2A and Supplementary Figure S2A, B) showed evident nucleosome arrays and were more similar to the observed nucleosome occupancy in *Zhang-MNase-invitro-salt* (black curves in Figure 2C and Supplementary Figure S2E, F). Interestingly, *Kaplan-MNase-invitro-salt* used 0.4:1 histone-to-DNA ratio in salt dialysis [3], while *Zhang-MNase-invitro-salt* used 1:1 [4], indicating that a higher histone-to-DNA ratio indeed results in a larger nucleosome number *in vitro*. We also note that the data in Figure 2 and Supplementary Figure S2 were standardized to have mean 0 and standard deviation 1, in order to facilitate the visual comparison of correlation; but, the outperformance of CEM was even more evident when normalization (dividing by the genome-wide mean) was used instead (Supplementary Figure S4, S5), in which case LM showed only a limited dynamic range in occupancy. Models involving only *GC* yielded similar comparison results, indicating that our conclusions did not depend on the specific sequence model employed (Supplementary Table S3, Supplementary Figures S6 and S7).

**Table 2::** Pearson and partial correlation analyses of CEM nucleosome occupancy predictions for three *in-vitro* datasets. *r*(data, prediction): Pearson correlation coefficients between observed and predicted nucleosome occupancy from *Model GC*, *GC*+*SR*, and *GC*+*SR*+*polyA*. The first row of partial correlation shows the partial correlation between observed and *Model GC*+*SR*-predicted nucleosome occupancy conditioned on *Model GC*-predicted nucleosome occupancy, representing any additional contribution from *SR*. The second row of partial correlation shows the partial correlation between observed and *Model GC*+*SR*+*polyA*-predicted nucleosome occupancy conditioned on *Model GC*+*SR*-predicted nucleosome occupancy, representing any additional contribution from *polyA*. All Pearson and partial correlation coefficients were calculated within “good regions” after filtering out outlier regions (Supplementary Methods, Section 1.5) and weighted by the region lengths in the reported average.

|  |  | Zhang-MNase-invitro-ACF | Kaplan-MNase-invitro-salt | Zhang-MNase-invitro-salt |
| --- | --- | --- | --- | --- |
| $r(\text{data}, \text{prediction})$ | *Model GC* | 0.775 | 0.729 | 0.589 |
|  | *Model GC+SR* | 0.775 | 0.730 | 0.592 |
|  | *Model GC+SR+polyA* | 0.775 | 0.755 | 0.606 |
| Partial correlation | *SR* | 0.048 | 0.036 | 0.081 |
|  | *polyA* | 0.018 | 0.287 | 0.161 |

Lastly, we note that the predicted nucleosome occupancy in Locke *et al.*’s manuscript did not show any clear nucleosome array in *Zhang-MNase-invitro-salt*, neither at the GAL10-1 locus (red curve in Supplementary Figure S1D of [16]) nor in the average profiles aligned at TSS/TTS (black dashed curves in Supplementary Figure S7B of [16]). By contrast, our implementations of LM did exhibit evident nucleosome arrays (red curve in Figure 2C and Supplementary Figure S2E,F). These discrepancies might have been caused by subtle differences in data processing affecting the normalization factor in Equation 1.

### 4.2 Contributions of G+C content, spatially resolved sequence motifs, and poly(dA:dT) tracts to *in-vitro* nucleosome occupancy

Diverse sequence features, such as *GC* [36], 10.5-bp periodicity [29, 37], and *polyA* [38], have been proposed to influence nucleosome landscape. However, which sequence feature actually contributes to which aspect of nucleosome landscape and the degree of these contributions are still under active debate. The flexibility of our CEM framework allows different sequence models to be studied on the same dataset (Equations 7, 8, and 9), providing an opportunity to dissect the complex relationship between different sequence features and different aspects of nucleosome landscape. This section quantifies how much the key sequence features that have received much attention in the community influence nucleosome occupancy.

We trained the aforementioned models in hierarchical order - *Models GC*, *GC*+*SR*, and *GC*+*SR*+*polyA* - on the three *in-vitro* nucleosome maps of *S. cerevisiae* studied in the previous section (Supplementary Table S4). To facilitate comparison between different datasets, we used a common template 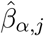 for all three datasets. This common template was constructed by combining average nucleotide frequencies of nucleosomes detected by MNase-seq [39] and chemical cleavage [40]; combining these two types of data help eliminate potential MNase digestion biases towards the ends of the nucleosome as well as chemical cleavage biases around the nucleosome dyad (Supplementary Figure S8). For all datasets considered, the three learned models had negative values of *μ*_C/G_ and positive values of *μ*_polyA_ (Supplementary Table S4), supporting the previous hypotheses that *GC* facilitates and *polyA* hinders nucleosome occupancy [36, 38]. To compare the relative contributions of the distinct sequence features to nucleosome occupancy, we calculated the Pearson correlation between the genome-wide observed and predicted nucleosome occupancy profiles (Table 2). *Model GC* achieved high correlation in all three datasets, and *Model GC*+*SR* showed essentially the same correlations as *Model GC*. These results suggest that nucleosome occupancy is predominantly determined by *GC*, while *SR*, including the 10.5-bp periodicity, does not play a significant role in determining nucleosome occupancy, consistent with previous studies [16, 29]. Furthermore, *Model GC*+*SR*+*polyA* showed slightly improved correlations in *Kaplan-MNase-invitro-salt* and *Zhang-MNase-invitro-salt*, but not in *Zhang-MNase-invitro-ACF*. To investigate whether *polyA* really does provide additional information about nucleosome occupancy on top of *GC* and *SR*, we calculated the partial correlation coefficient between the observed and *Model GC*+*SR*+*polyA* predicted nucleosome occupancy conditioned on the *Model GC*+*SR* prediction (Table 2) (Supplementary Methods, Section 1.6) [41, 42]. The calculated partial correlation was around 0.2 in both *Kaplan-MNase-invitro-salt* and *Zhang-MNase-invitro-salt*, but negligible in *Zhang-MNase-invitro-ACF*. By contrast, similar partial correlations for *Model GC*+*SR* conditioned on *Model GC* was < 0.1 for all three datasets (Table 2). Therefore, for *in-vitro* nucleosome maps obtained from salt dialysis, *polyA* does make a small contribution to determining genome-wide nucleosome occupancy, while the contribution from *SR* is negligible.

**Figure 3::**
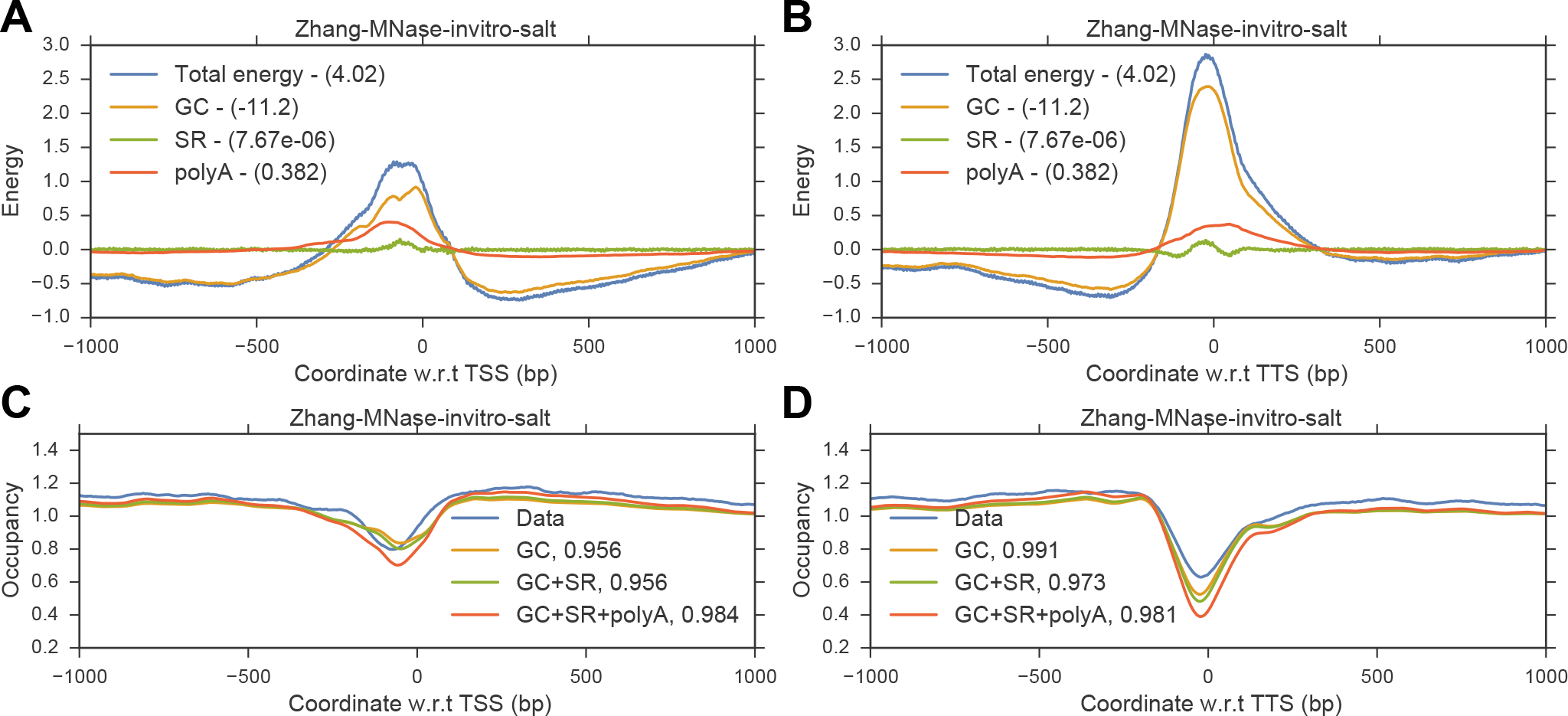
Contributions from *GC*, *SR*, and *polyA* in shaping nucleosome occupancy at TSS and TTS. (A) Nucleosome-positioning energy in *Model GC*+*SR*+*polyA* attributable to *GC* (yellow curve), *SR* (green curve), and *polyA* (red curve) aligned and averaged at TSS of all genes. The total energy is shown in blue. Each energy component is subtracted by its genome-wide mean shown in the legends to facilitate visualization. (B) Same as (A), but aligned at TTS. (C) Observed (blue curve) and predicted nucleosome occupancy from *Models GC* (yellow curve), *GC*+*SR* (green curve), and *GC*+*SR*+*polyA* (red curve), aligned and averaged at TSS and normalized by the genome-wide mean. Pearson correlation coefficients between observation and prediction are shown in the legends. (D) Same as (C), but aligned at TTS.

Locally, it is well known that nucleosome occupancy tends to be reduced around TSS and TTS, resulting in nucleosome depleted regions (NDR) [31]. To study which sequence features predominantly contribute to the formation of NDR, we plotted the nucleosome-positioning energies at TSS and TTS from the three separate components of *GC*, *SR*, and *polyA* in the learned *Model GC*+*SR*+*polyA* (Figure 3 and Supplementary Figure S9). Again, *GC* was the primary component responsible for the energy barrier at both TSS and TTS (Figure 3A,B and Supplementary Figure S9A,B,E,F). compared with *GC*, *polyA* showed detectable but smaller energy barriers in datasets obtained from salt dialysis (Figure 3A,B and Supplementary Figure S9A,B), but not in *Zhang-MNase-invitro-ACF* (Supplementary Figure S9E,F). In comparison, *SR* showed much suppressed energy barriers. Interestingly, CEM predicted higher energy barriers and lower nucleosome occupancies at TTS than at TSS in all three datasets, consistent with the observed nucleosome occupancy. Finally, unlike in datasets obtained from salt dialysis, neither *polyA* nor *SR* showed any influence on nucleosome occupancy in *Zhang-MNase-invitro-ACF*, suggesting that chromatin assembly factors such as ACF might overwrite certain intrinsic sequence preferences of nucleosome formation.

### 4.3 The dependence of MNase-derived nucleosome occupancy maps on G+C content is substantially biased by MNase digestion

We have shown in the previous section that *GC* is the primary determinant of the nucleosome occupancy derived from MNase-seq experiments. However, MNase has a substantial *GC* bias in its cleavage preference [27, 28]. To further investigate the *GC* dependence of nucleosome occupancy, we calculated the Pearson correlation between *GC*, *polyA*, and observed nucleosome occupancy generated by an independent chemical cleavage method that did not involve MNase (*Brogaard-chemical-invivo-WT*) [40] (Figure 4A). Since the chemical cleavage data were obtained *in vivo*, we analyzed an *in-vivo* dataset generated by MNase-seq (*Kaplan-MNase-invivo-WT*) [3] for comparison (Figure 4D). As before, chemical-cleavage-derived occupancy showed positive correlation with *GC* and negative correlation with *polyA* (Figure 4A). However, unlike MNase-derived occupancy which correlated, either positively or negatively, most strongly with *GC* (Figure 4D), we found that chemical-cleavage-derived occupancy correlated most strongly with *polyA* (Figure 4A). Furthermore, partial correlation analysis showed that there was little direct correlation between *GC* and chemical-cleavage-derived occupancy after conditioning on *polyA* (Figure 4B), while the correlation between *polyA* and chemical-cleavage-derived occupancy was mostly retained even after conditioning on *GC* (Figure 4B). This result shows that the marginal correlation between *GC* and chemical-cleavage-derived occupancy was largely a secondary consequence of the negative correlation of both variables with *polyA*. By contrast, the *GC* dependence of MNase-derived nucleosome occupancy was almost unchanged after conditioning on *polyA* (Figure 4E). Other independent MNase-derived *in-vivo* and *in-vitro* datasets showed similar trends (Supplementary Figure S10). Therefore, the *GC* dependence of nucleosome occupancy was specific to MNase-derived datasets, indicating that this *GC* dependence might be an artifact of MNase digestion biases.

To further resolve the experiment-specific relationship between nucleosome occupancy, *GC*, and *polyA*, we divided the genome into 1000-bp segments and binned the segments according to their *GC* and poly(dA:dT) content. We then plotted the average nucleosome occupancy in the bins as a heat map (Figure 4C for chemical cleavage occupancy, Figure 4F for MNase). The existence of a left-to-right gradient in Figure 4C demonstrated the direct correlation between *polyA* and chemical-cleavage-derived nucleosome occupancy, while the lack of a top-to-bottom gradient in Figure 4C showed that there was little direct correlation between *GC* and chemical-cleavage-derived nucleosome occupancy. This figure also confirmed that it was the negative correlation between *GC* and *polyA* that gave rise to the spurious correlation between *GC* and chemical-cleavage-derived nucleosome occupancy. By contrast, for the MNase-derived dataset, both *GC* and *polyA* showed direct correlation with nucleosome occupancy, illustrated by the existence of both weaker left-to-right and stronger top-to-bottom gradient in Figure 4F. Analysis of other independent MNase-derived datasets yielded similar results (Supplementary Figure S10). These results thus further confirmed that the *GC* dependence of nucleosome occupancy was specific to MNase-derived datasets.

We reasoned that if the *GC* dependence of MNase-derived nucleosome occupancy was truly a consequence of MNase digestion biases, then the correlation between *GC* and nucleosome occupancy should depend on the MNase digestion level. That is, the higher the digestion level, the stronger should be the correlation between *GC* and nucleosome occupancy. To test this hypothesis, we analyzed a dataset where MNase digestion was performed for 2.5, 5, 10, and 20 minutes [43] (Figure 4G, H). As expected, the correlation between *GC* and nucleosome occupancy increased with digestion time (Figure 4G), as did the partial correlation between *GC* and occupancy conditioned on *polyA* (Figure 4H).

**Figurer 4:**
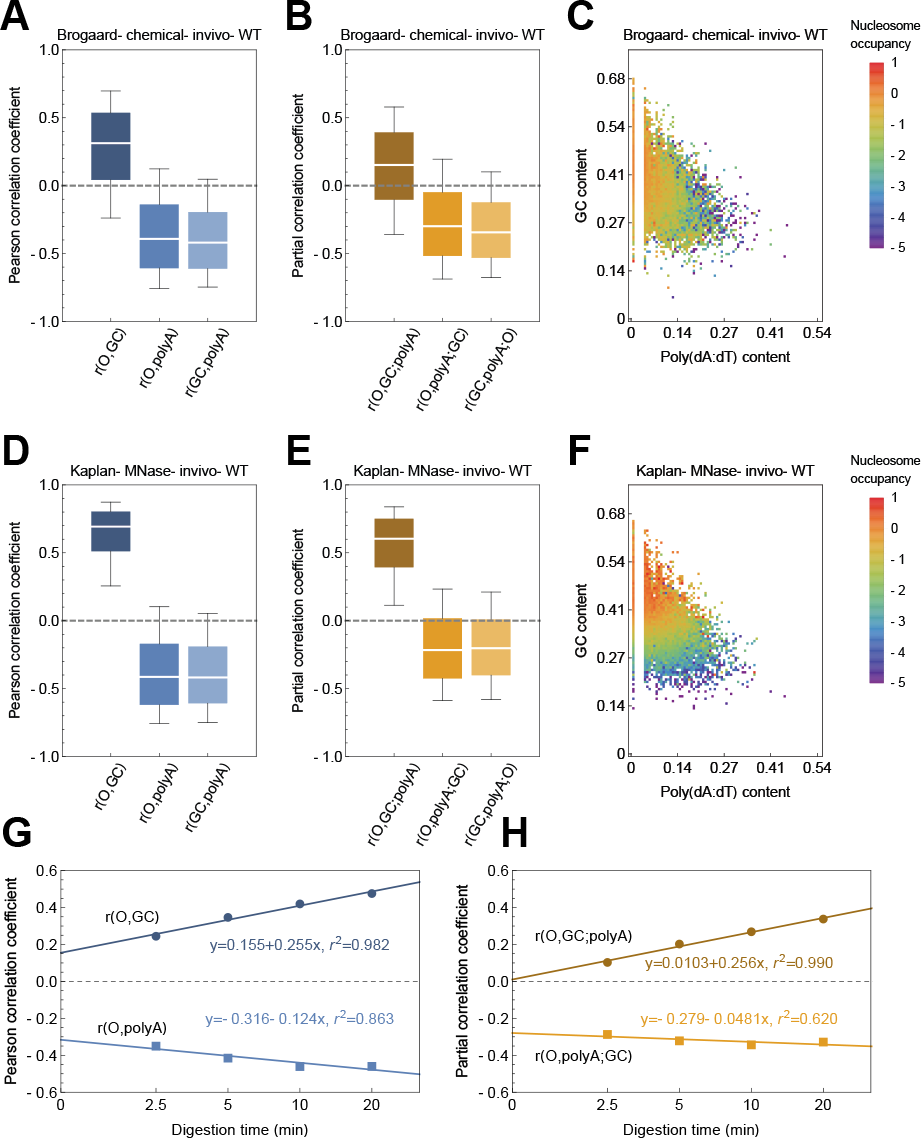
The dependence of MNase-derived nucleosome occupancy on G+C content is substantially biased by MNase digestion. (A) Distribution of pairwise Pearson correlation coefficients between chemical-cleavage-derived nucleosome occupancy, *GC*, and *polyA*, calculated on 1000-bp intervals tiling the “good regions.” (B) Distribution of pairwise partial correlation coefficients between chemical-cleavage-derived nucleosome occupancy, *GC*, and *polyA*, conditioning on the third variable, calculated on 1000-bp intervals tiling the “good regions.” (C) A heatmap of nucleosome occupancy as a function of *GC* and *polyA*. The *S. cerevisiae* genome was divided into 1000-bp segments, and each segment was then assigned to a 2-dimensional bin of given GC and poly(dA:dT) content. Color indicates the average nucleosome occupancy in each bin. (D-F) Same as (A-C), but for MNase-derived nucleosome occupancy. (G) Median of the Pearson correlation coefficients of *GC* and *polyA* with MNase-derived nucleosome occupancy at different digestion levels, calculated on 1000-bp intervals tiling the “good regions.” Linear extrapolation was performed to infer the correlation coefficient at digestion time 0. (H) Same as (G), but for partial correlation coefficients.

Interestingly, the Pearson and partial correlation coefficients showed an almost linear relation with the logarithm of digestion time, although there were only four data points. This linear relationship allowed us to extrapolate the correlation coefficients at digestion time 0, which could be thought of as representing the true biological condition without any MNase bias. Strikingly, the partial correlation between *GC* and nucleosome occupancy almost dropped to 0 at digestion time 0 in this extrapolation analysis (Figure 4H), suggesting that almost all of the observed *GC* dependence of MNase-derived nucleosome occupancy was due to MNase digestion biases. By contrast, the partial correlation between *polyA* and nucleosome occupancy conditioned on *GC* changed little with digestion time and was largely retained at the extrapolated time 0 (Figure 4H), further supporting that the *polyA* dependence of nucleosome occupancy reflected a true biological signal.

### 4.4 Contributions of G+C content, spatially resolved sequence motif, and poly(dA:dT) tracts to *in-vitro* nucleosome positioning

We now turn to investigating the sequence determinants of translational and rotational nucleosome positioning. We trained the *Models GC*, *GC*+*SR*, and *GC*+*SR*+*polyA* on *in-vitro* nucleosome maps generated through MNase-seq experiments (Supplementary Table S4) and predicted nucleosomes genome-wide by progressively calling genomic locations with the largest predicted *n*_1_ as nucleosome start positions, while constraining that the called nucleosomes cannot overlap with each other (Supplementary Methods, Section 1.5). The nucleosome calls were thus ranked by the value of *n*_1_. Similarly, positioned nucleosomes were also identified from experimental data by using the same algorithm. To assess the agreement between observed and predicted nucleosome positions, we examined top *N* nucleosomes from the respective ranked lists, where *N* was set to be the nucleosome number predicted by *Model GC* in each dataset (Supplementary Table S4; Supplementary Methods, Section 1.5). The three models showed modest and similar prediction power on single nucleosome positions (Figure 5A), being able to predict about 50% of observed nucleosomes within 50 bps. In particular, adding the *SR* component to nucleosome-positioning energy did not significantly improve the prediction accuracy, agreeing with our previous finding based on spectral analysis that sequence periodicity does not play a major role in translational positioning [29].

**Figure 5:**
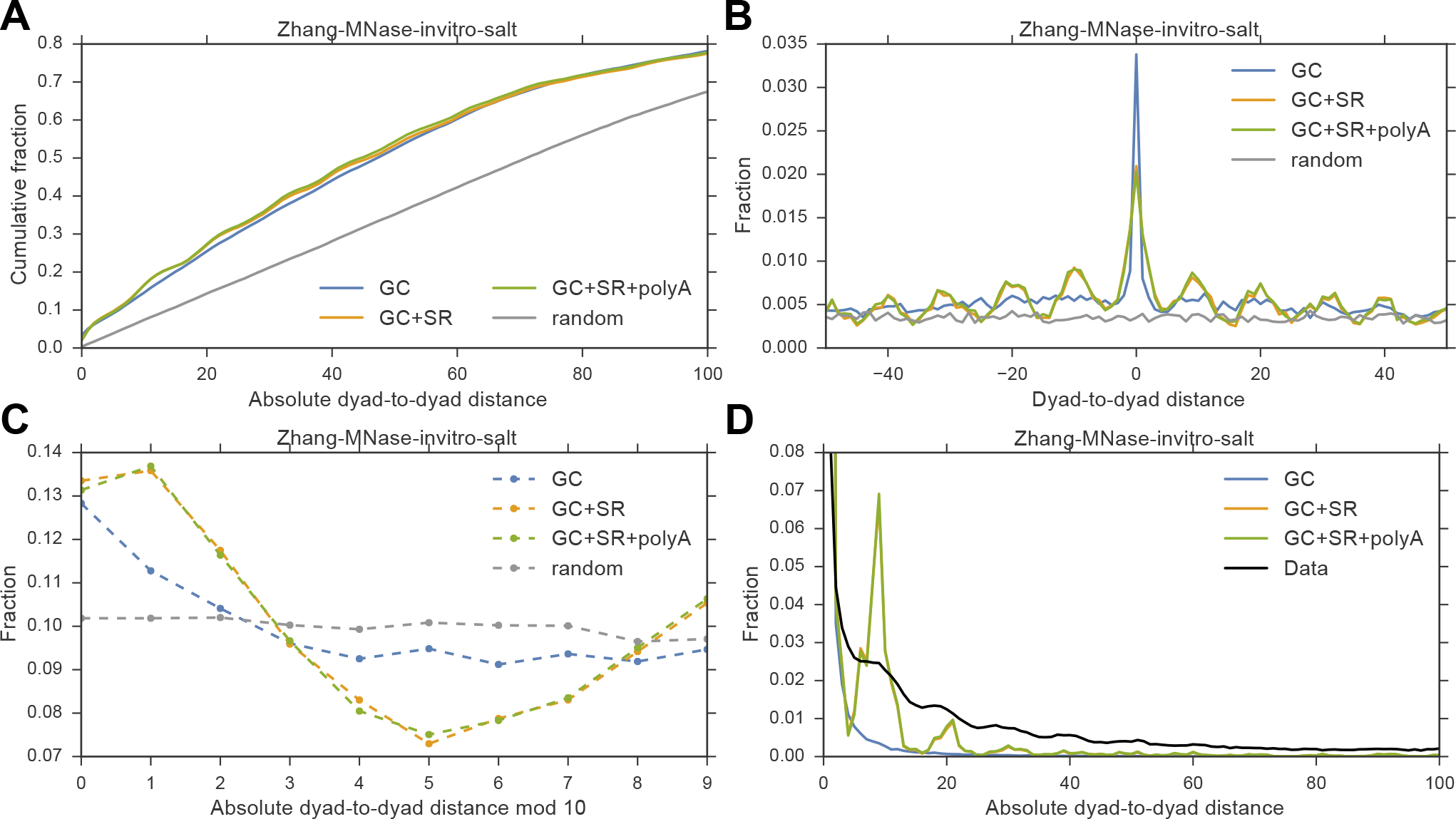
Spatially-resolved sequence motifs, including the 10.5-bp periodicity, facilitate the rotational but not translational positioning of nucleosomes. (A) cumulative distribution of absolute dyad-to-dyad distance between predicted and observed nucleosome positions. Grey curve represents a random control using non-overlapping uniformly distributed nucleosomes. (B) Distribution of dyad-to-dyad distance between predicted and observed nucleosome positions. (C) Distribution of absolute dyad-to-dyad distance between predicted and observed nucleosome positions mod 10 bp. (D) Distribution of dyad-to-dyad distance between redundant nucleosomes.

By contrast, the distribution of dyad-to-dyad distance between the nucleosome positions observed in sequencing data and those predicted by the models including an *SR* component was enriched for multiples of 10 bp (Figure 5B), and the rotational nucleosome positions were also better predicted by these models (Figure 5C). In order to better capture cell-to-cell variation in nucleosome positions, we progressively identified a redundant set of nucleosomes containing five times the above number of nucleosomes in each dataset. The dyad-to-dyad distance between nucleosomes within this redundant set exhibited a 10-bp oscillatory pattern emerging from rotational positioning of nucleosomes (black curve in Figure 5D). Importantly, this 10-bp oscillation could be recapitulated by a model only if it included the *SR* component in its nucleosome-positioning energy (Figure 5D). These results thus demonstrated that spatially-resolved sequence motifs, including the 10.5-bp periodicity, facilitated the rotational positioning of nucleosomes, again confirming our previous conclusion based on spectral analysis [29]. Analysis of other *in-vitro* nucleosome maps provided similar results, except that *Zhang-MNase-invitro-ACF* showed little sequence dependence other than *GC*, as previously mentioned (Supplementary Figure S11).

### 4.5 Models on *in-vivo* nucleosome maps exhibit a high degree of experiment-specific variability

We have so far focused on *in-vitro* nucleosome maps, since *in-vitro* conditions provide a simplified platform for studying intrinsic sequence preferences hidden in the nucleosome landscape. *In vivo*, DNA sequence is only one of many factors that can influence nucleosome positioning. *Trans*-factors, such as chromatin remodelers, transcription factors and RNA polymerase, can actively regulate nucleosome positions. In order to quantify the role of DNA sequence in determining the *in-vivo* nucleosome landscape, we applied CEM to available *in-vivo* nucleosome maps. See Supplementary Methods, Section 1.4 for detailed information about these datasets and our labeling convention. CEM-predicted nucleosome occupancy achieved consistently higher correlation with observed nucleosome occupancy than the LM prediction (Supplementary Table S5), demonstrating that the benefits of CEM generalized to *in-vivo* datasets. However, CEM trained on different *in-vivo* nucleosome maps yielded discordant results (Supplementary Table S5). In particular, the estimated nucleosome number, although consistently larger than that predicted by LM, varied between ~36000 and ~56000.

Since all curated *in-vivo* nucleosome maps were generated using wild-type *S. cerevisiae* in log phase, one would expect similar nucleosome numbers in these datasets. Indeed, a quantitative estimation of nucleosome number by matching the Fourier transform of observed nucleosome occupancy profiles showed that all analyzed *in-vivo* nucleosome maps contained a similar number of ~65000 nucleosomes (Supplementary Figure S12), suggesting that CEM was consistently underestimating the nucleosome number in these datasets (Supplementary Table S5). Visually comparing predicted and observed nucleosome occupancy profiles showed that, for datasets where nucleosome number was significantly underestimated (*Kaplan-MNase-invivo-WT* and *Ocampo-MNase-invivo-WT*), CEM-predicted nucleosome occupancy was indeed missing a large fraction of nucleosomes evident from experimental data (Supplementary Figure S13). Of note, the observed nucleosome occupancy in these datasets showed peculiar long-range fluctuations which were not observed in *McKnight2016-MNase-invivo-WT-log-80* and *McKnight2015-MNase-invivo-WT-log-50*, where nucleosome number was only modestly underestimated (Supplementary Figure S13). Interestingly, the latter two datasets, *McKnight2016-MNase-invivo-WT-log-80* and *McKnight2015-MNase-invivo-WT-log-50*, that we were able to model better, were from the same lab and generated by the same author(s). Fourier analysis of the observed nucleosome occupancy further demonstrated that these two datasets also had relatively large spectral density at the peak around ~200 bp corresponding to a nucleosome-size signal. By contrast, other *in-vivo* nucleosome maps contained smaller spectral density at the peak around ~200 bp, but had more spectral energy in long-range components (Supplementary Figure S12A). These results suggested that experiment-specific noises and biases, such as a trending effect, might be confounding the CEM optimization.

Since the two datasets *McKnight2016-MNase-invivo-WT-log-80* and *McKnight2015-MNase-invivo-WT-log-50* exhibited the smallest long-range trending effect and resulted in only a modest underestimation of nucleosome number, we used these two datasets to study the *in-vivo* nucleosome landscape by training the aforementioned *Models GC*, *GC*+*SR*, and *GC*+*SR*+*polyA* (Supplementary Table S6, S7). As in the *in-vitro* case, *GC* was still the primary factor determining *in-vivo* nucleosome occupancy (Supplementary Table S6), although this *GC* dependence might have arisen from MNase digestion biases, as previously described. Also similar to the *in-vitro* findings, *polyA* made a small contribution to determining *in-vivo* nucleosome occupancy (Supplementary Table S6). One difference was that *SR* seemed to have a greater impact on nucleosome occupancy *in vivo* than *in vitro* (Supplementary Table S6), especially at the TSS (Supplementary Figure S14). For single nucleosome positions, the prediction accuracy based on sequence information alone was very modest, although *SR* still showed signatures of facilitating rotational positioning (Supplementary Figure S15).

### 4.6 *In-vivo* nucleosome occupancy around TSS is partially determined by DNA sequence

It has been known for a long time that nucleosome occupancy *in vivo* tends to be reduced around TSS, leading to a nucleosome depleted region (NDR), and that the average nucleosome occupancy downstream of TSS exhibits an oscillatory pattern reflecting statistical positioning [18, 31]. It has been argued that DNA sequence itself is not sufficient to explain the NDR and oscillatory pattern [16] and that active remodeling by *trans*-factors are needed [23]. Ostensibly consistent with these claims, the CEM-predicted occupancy using *Model GC*+*SR*+*polyA*, averaged over all genes at TSS, did not show an NDR that was as deep as in the observed occupancy (Figure 6A, Supplementary Figure S16A). Furthermore, the oscillations in CEM-predicted average occupancy were very weak (Figure 6A, Supplementary Figure S16A), seemingly supporting the idea that DNA sequence alone is not able to fully explain the energy barrier present at TSS.

**Figure 6:**
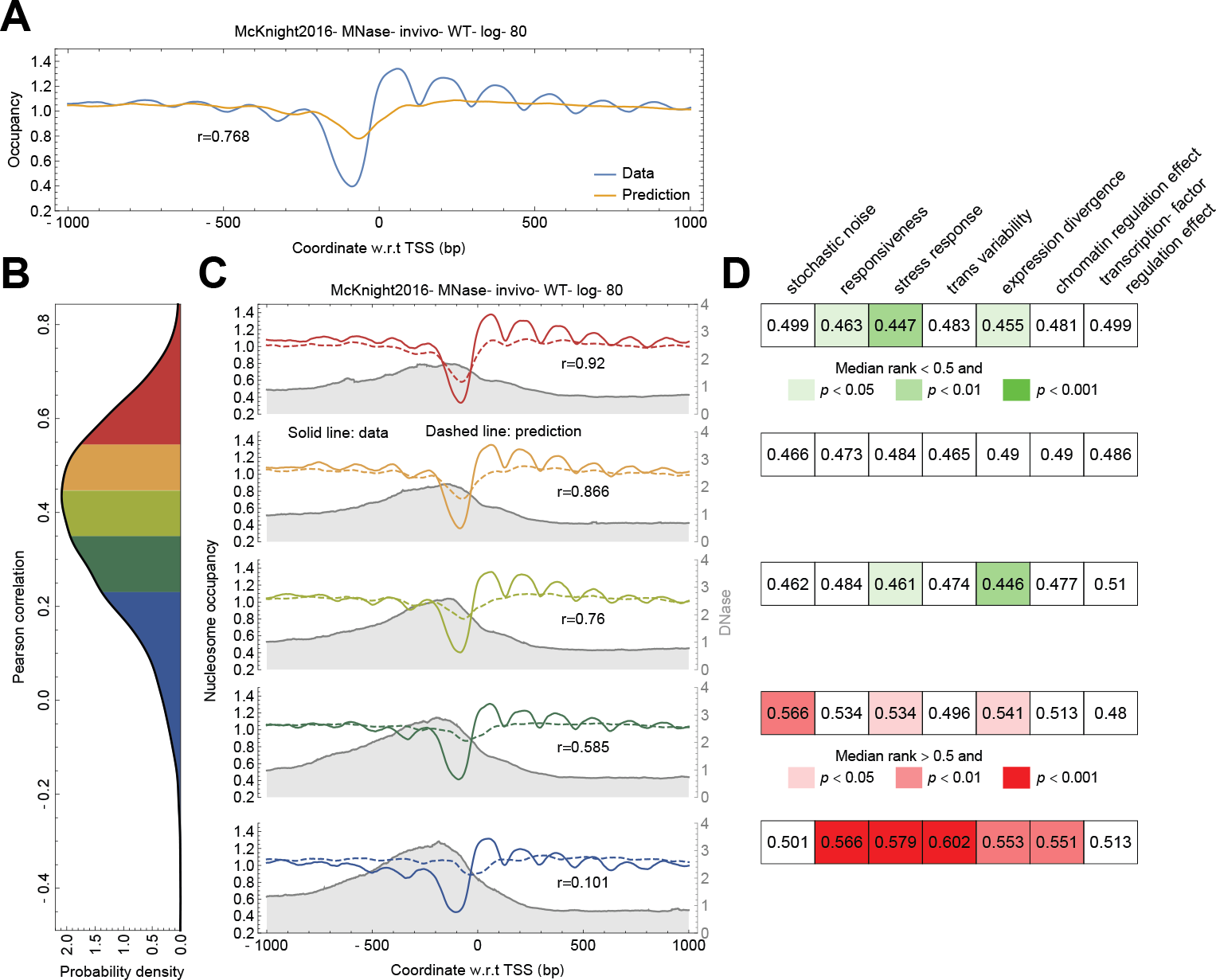
*In-vivo* nucleosome occupancy around TSS is partially determined by DNA sequence. Shown are the results using the *Model GC*+*SR*+*polyA* trained on *McKnight2016-MNase-invivo-WT-log-80*. (A) Observed (blue) and predicted (yellow) nucleosome occupancy aligned at TSS and averaged over all genes. (B) Genes were ranked by the Pearson correlation coefficient between observed and predicted nucleosome occupancy within ±1 kb of TSS and divided into quintiles (different colors). The distribution of these correlation coefficients are shown. (C) Observed (solid curves) and predicted (dashed curves) nucleosome occupancy aligned at TSS and averaged over the genes within each quintile from (B). Average DNase I hypersensitivity [44] (gray curve and shade) is also shown for genes within each quintile. (D) Enrichment analysis for expression variability [45]. All genes are ranked between 0 and 1 according to a variability measure. The median rank of genes within each quintile is shown. A permutation test is performed to assess whether the genes within each quintile are significantly enriched for high or low variability.

To investigate the nucleosome occupancy around TSS in greater detail, however, we ranked the genes according to their predictability, as measured by the Pearson correlation between observed and predicted nucleosome occupancy within ±1 kb from TSS, and divided the genes into quintiles (Figure 6B, Supplementary Figure S16B). For genes in the fifth quintile showing the highest predictability, the NDR depth and oscillatory pattern in predicted occupancy better matched the observed patterns (Figure 6C, Supplementary Figure S16C); for this subset of genes, DNA sequence thus played a more significant role in regulating nucleosome occupancy at TSS. Interestingly, this same subset of genes showed less DNase hypersensitivity [44] around TSS (Figure 6C, Supplementary Figure S16C), smaller expression variability as measured in [45] (Figure 6D, Supplementary Figure S16D), and a slightly lower histone turnover rate [46] (Supplementary Figure S17). Importantly, the oscillation of average nucleosome occupancy for this class of genes was predicted from DNA sequence alone, using the simplest model accounting for only hard-core interactions between adjacent nucleosomes. In sharp contrast, previous studies either failed to reproduce this oscillatory pattern using sequence alone [16], potentially due to their underestimation of nucleosome number, or required artificially constructed energy barriers at TSS [23]. The fact that DNA sequence alone could partially recapitulate the average nucleosome occupancy in a subset of genes highlighted that different genes might utilize distinct mechanisms to regulate nucleosome formation at TSS.

### 4.7 *In-vivo* nucleosome occupancy around TTS is primarily determined by DNA sequence

Finally, we investigated the pattern of *in-vivo* nucleosome occupancy around TTS. Unlike at TSS, the *Model GC*+*SR*+*polyA* was able to fully capture the nucleosome depletion at TTS (Figure 7A, Supplementary Figure S18A). The slight discrepancy between prediction and observation at TTS could be explained by the existence of a nearby TSS [47]. When sorted by the distance to the closest TSS (Figure 7B, Supplementary Figure S18B), the TTS isolated from any TSS had an even better agreement between predicted and observed nucleosome occupancy (Figure 7C, Supplementary Figure S18C); by contrast, those TTS close to TSS showed greater deviation from prediction (Figure 7C, Supplementary Figure S18C). We note, however, that these observations might be specific to MNase-derived nucleosome maps. For instance, the sharper energy barrier at TTS predicted by CEM was mostly attributable to *GC* (Supplementary Figure S14), but according to our previous analysis, the *GC* dependence of MNase-derived nucleosome maps were subject to substantial MNase digestion biases. Moreover, nucleosome maps obtained from experimental methods avoiding MNase digestion showed much less pronounced depletion at TTS than the maps obtained from MNase-seq [48].

## 5 Discussion

In this paper, we have developed a cross-entropy method (CEM) for simultaneously learning the nucleosome number and nucleosome-positioning energy from empirical genome-wide nucleosome maps. CEM circumvents a crucial computational difficulty encountered by previous approaches when estimating the nucleosome number [16]. Applying our framework to various *in-vitro* and *in-vivo* datasets in *S. cerevisiae* has provided a comprehensive quantification of the role of DNA sequence in shaping the genome-wide nucleosome landscape that includes the distinct concepts of nucleosome occupancy and nucleosome positioning.

Using *in-vitro* nucleosome maps, we have shown that spatially resolved sequence patterns, including the controversial 10.5-bp nucleotide periodicity, facilitate rotational, but not translational positioning, consistent with the previous conclusion based on categorical spectral analysis of nucleosomal DNA [29]. In our models, *GC* is the primary determinant of MNase-derived nucleosome occupancy, while the ostensibly minor role of *polyA* may be under-estimated due to a confounding effect. More precisely, our partial correlation analysis has demonstrated that the *GC* dependence of MNase-derived nucleosome occupancy may be largely attributable to MNase digestion biases, in line with a recent deep-learning study showing that *GC* contributes to the classification of only a small fraction of nucleosomal sequences in *S.cerevisiae* [49]. However, we cannot rule out the possibility that the intrinsic sequence preferences of MNase digestion and nucleosome formation are partially overlapping. Using currently available sequencing data, it is difficult to dissect how much of the *GC* dependence is due to MNase bias and how much, if any, is truly nucleosome preference. Further *in-vitro* studies using MNase-independent experiments, such as the chemical cleavage method [40, 50, 51], should improve our understanding in the future. By contrast, the effect of *polyA* on nucleosome occupancy is robust and independent of the specific experimental method used, suggesting that *polyA*, together with the steric exclusion between adjacent nucleosomes, may be the truly dominant factor in determining genome-wide nucleosome occupancy.

**Figure 7:**
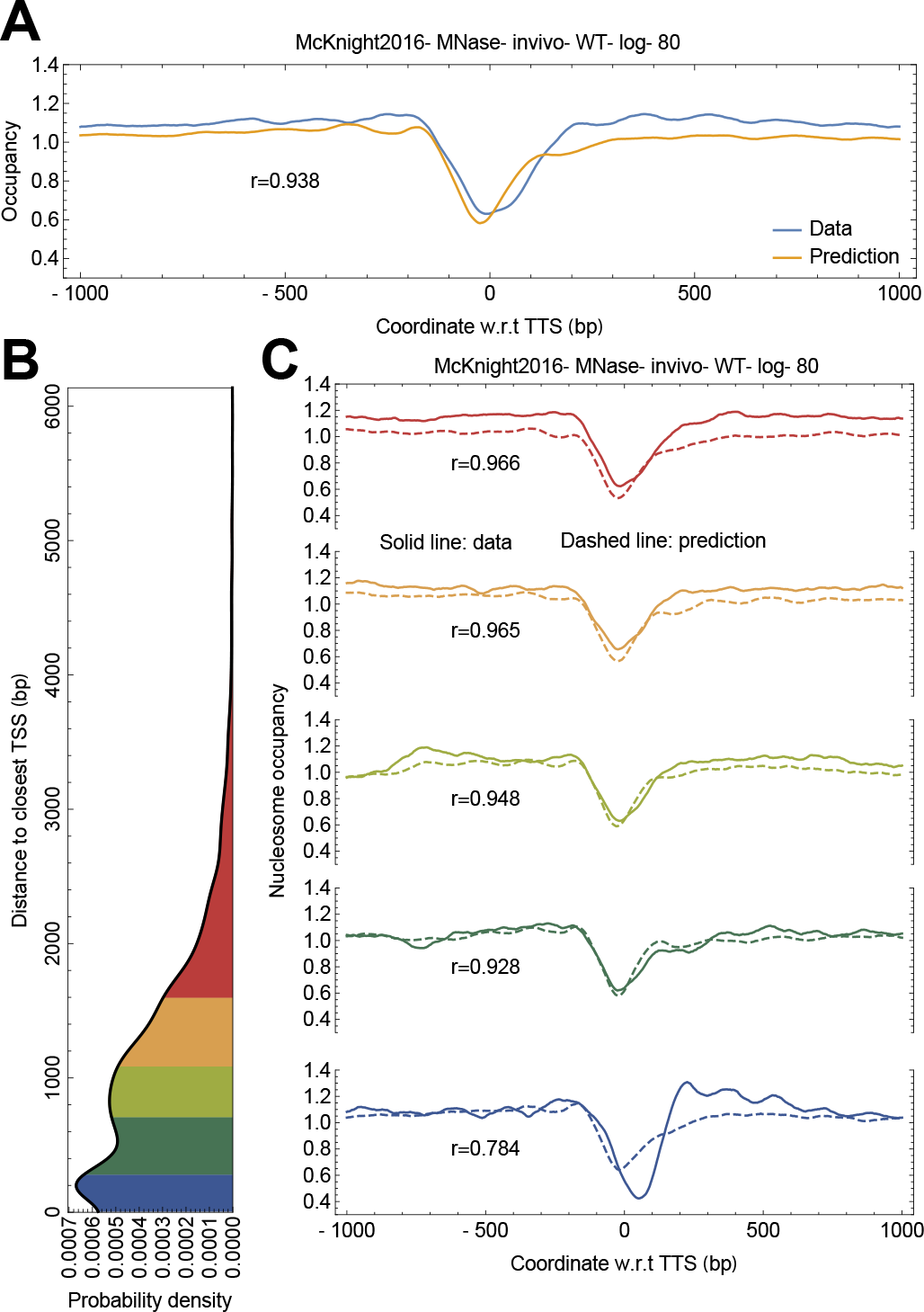
*In-vivo* nucleosome occupancy around TTS is primarily determined by DNA sequence. Shown are the results using the *Model GC*+*SR*+*polyA* trained on *McKnight2016-MNase-invivo-WT-log-80.* (A) Observed (blue) and predicted (yellow) nucleosome occupancy aligned at TTS and averaged over all genes. (B) Genes are ranked by the distance from their TTS to the closest TSS and divided into quintiles (different colors). The distribution of these distances are shown. (C) Observed (solid curves) and predicted (dashed curves) nucleosome occupancy aligned at TTS and averaged over the genes within each quintile from (B).

Applying CEM to *in-vivo* nucleosome maps has shown, for the first time, that the canonical oscillatory pattern in average nucleosome occupancy around TSS can be partially recapitulated by DNA sequence and steric exclusion of nucleosomes for a subset of genes. We believe that previous studies have not been able to capture this pattern, because of the aforementioned underestimation of true nucleosome number [16]. This result also highlights a key difference between *in-vitro* and *in-vivo* nucleosome maps: *in-vivo* conditions tend to yield a larger nucleosome number, or equivalently, higher chemical potential. Although some *in-vitro* experiments have used histone-to-DNA ratios close to those under *in-vivo* conditions [4], histone chaperones and chromatin remodelers may actively package histones onto DNA, effectively increasing the chemical potential *in vivo*. This difference between *in-vitro* and *in-vivo* conditions, although previously noted [15, 22], has not been fully appreciated to date, especially regarding the formation of regularly spaced nucleosome arrays around TSS.

Specifically, the oscillatory pattern in average nucleosome occupancy around TSS has been thought to be a unique feature of the *in-vivo* nucleosome landscape created by regulatory factors other than DNA sequence itself. However, the fact that DNA sequence alone in our *in-vivo* models can reproduce this oscillatory pattern for select genes suggests that, given high enough chemical potential, this pattern may be also produced *in vitro*. This hypothesis may be tested by mapping nucleosome positions *in vitro* with an even higher histone-to-DNA ratio than that used in [4]. In fact, our analysis shows that nucleosome arrays are already vaguely visible in some *in-vitro* nucleosome maps [4] (Figure 2C, Supplementary Figure S5E,F) and that tuning up the chemical potential to match the expected *in-vivo* nucleosome number indeed creates a more pronounced oscillatory pattern around TSS (Supplementary Figure S19). We also note that the effective chemical potential may be different at different genomic locations, especially *in vivo* [15]. A future study where separate models are trained on distinct genomic segments may help elucidate our hypothesis [15].

Our method represents a unified framework that accounts for both intrinsic sequence preference of nucleosome formation and steric exclusion between neighboring nucleosomes. In order to fully complete the story, other factors such as chromatin remodelers, transcription factors, and RNA polymerase need to be taken into account. One advantage of our computational framework based on CEM is that it can be potentially extended to include these regulatory factors. To a first order approximation, the presence of these *trans*-factors can be considered as modulating the effective nucleosome-positioning energy in addition to DNA sequence. For example, an additional term that linearly depends on the ChIP-seq intensity of relevant *trans*-factors can be introduced into Equation 9. Another related strategy is to use CEM to study specific functions of individual chromatin remodelers, utilizing the data generated through *in-vitro* genome-wide reconstitution of nucleosomes in the presence of individual chromatin remodelers [52]. For example, it has been shown that RSC clears the promoter by reading poly(dA:dT) as a directional nucleosome removal signal [52]. We expect this effect to be captured by a relative change in μ_polyA_ in Equation 9. Similarly, INO80 has been shown to help position the +1 nucleosome [52], suggesting that INO80 may act as an additional energy barrier at TSS. Thus, incorporation of INO80 chIP-seq information into the sequence model may help improve the prediction of nucleosome landscape. Lastly, it is known that ISW1 can tighten the spacing in nucleosome arrays downstream of TSS [52]. A plausible hypothesis is that ISW1 increases the effective chemical potential so that the nucleosomes in the region are more tightly packed. All of these aspects can potentially be captured by CEM with different energy models.

Finally, it is important to note that other statistical mechanics-inspired models that relax the assumption of hard-core interactions have been also used to study nucleosomes. For example, nucleosome unwrapping and breathing can be captured by a soft-core interaction [25] or variable nucleosome size [20, 26]. More complex nearest neighbor interactions imposed by higher-order chromatin structure can be modeled [23, 24]. Models beyond equilibrium statistical mechanics have also been attempted [53–55]. Nevertheless, this paper demonstrates that even the simplest model involving only steric exclusion can still provide new insights into the principles that guide nucleosome occupancy and nucleosome positioning.

